# ScanFold3: a robust and rapid approach for identifying structured RNA targets in genomes and transcriptomes

**DOI:** 10.64898/2026.10.29.762533

**Authors:** Evelyn C. Coppenbarger, Abdelraouf O. Dapour, Ryan J. Andrews, Walter N. Moss

**Affiliations:** The Roy J Carver Department of Biochemistry, Biophysics and Molecular Biology, Iowa State University, Ames, IA, United States; Bioinformatics and Computational Biology Program, Iowa State University, Ames, IA, United States; Department of Biochemistry, University of Utah, Salt Lake City, UT, United States

## Abstract

**Background:** The rapid expansion of high-throughput transcriptomic datasets has created a critical need for scalable tools to identify functional RNA secondary structures. While thermodynamics-based folding is a well-established method, most algorithms lack the scalability required for genomic-scale datasets due to O(n^3^) time complexity.

**Methods:** We present ScanFold3, an enhanced version of the ScanFold pipeline that optimizes the “Fold” step for modern computing needs. This version models consensus structure generation as a minimum perfect weighted matching problem on a graph. To ensure maximum scalability, we implemented a Greedy Approximation Algorithm and optimized performance-critical components in C++.

**Results:** Benchmarking on sequences up to 128 kb demonstrates that ScanFold3 achieves a 38-fold to 129-fold increase in speed over ScanFold2. Validation against annotated human RNAs shows that ScanFold3 offers improved positive predictive value (PPV). Most significantly, when ScanFold3 outputs are used as structural constraints for global refolding, it outperforms previous versions across all metrics, including the F1 score.

**Conclusions:** ScanFold3 provides a robust, speedy, and user-friendly method for characterizing the thermodynamics of novel RNA and DNA sequences. It is designed as an efficient first step for high throughput bioinformatic pipelines, capable of paring down chromosome-scale transcriptomic data into stable structures amenable to advanced analysis.

## Introduction

RNA structures have been implicated in processes a number of diverse biological processes, such as gene regulation Morris and Mattick (2014), viral frameshifting (Giedroc, Theimer et al. 2000), and RNA splicing (Mattick and Makunin 2006). The formation of these structures is highly-dependent on their thermodynamic stability. RNA stability is driven by base stacking interactions and hydrogen bonding between the edges of bases which also confers specificity (Mathews, Sabina et al. 1999, Mathews, Moss et al. 2010, Turner and Mathews 2010). The pattern of base pairs— the “2D” or “secondary” structure—constrains energetically favorable “3D” or “tertiary” structures folds. Because base pairing is generally predictable based on the sequence and known thermodynamic properties of nitrogenous bases, secondary structure prediction remains a cornerstone of RNA research.

RNA is able to form helical structures similar to DNA, though unlike DNA these structures are formed within individual strands and typically have unpaired (bulged or looped) nucleotides. The result is a structure, with a mix of paired and unpaired nucleotides: e.g., “hairpin loops” where the RNA strand bends back on itself, terminating in a loop structure. RNA structures can be decomposed into sub-structures: stacking dinucleotides, where two pairs are adjacent and able to stabilize each other via stacking interactions; bulge loops, where a stacking dinucleotide pair is broken by insertion of an unpaired nucleotide on one side; internal loops, where a stacking interaction is broken by inserting an unpaired nucleotide on both sides; hairpin loops, also called terminal loops, where the molecule has folded back on itself and thus the loop is bound by only one dinucleotide, rather than two; and multi-branch loops, where more than one stem-loops branch off of a loop.

One of the most common methods for predicting RNA secondary structures due to its speed and reliability is the Zuker Algorithm (Zuker and Stiegler 1981), a dynamic programming based approach which builds up a matrix of possible base pairs where each entry is the free energy of the most favorable substructure found between those pairs. Each structure can be decomposed down into substructures until one reaches the level of individual stacking energies and (generally) penalties for being unpaired. The energetic parameters of these structural motifs can be empirically measured and many have been (Mathews, Sabina et al. 1999, Turner and Mathews 2010). The matrix is built up, then a traceback algorithm works backwards to find each pair involved in the best substructure until a final minimum free energy (MFE) structure is found (Eddy 2004).

While this is a powerful approach, even the optimized form of the algorithm has a time complexity of O(n^3^) with respect to sequence length (Lyngsø, Zuker et al. 1999), which limits its scalability. Beyond computational expense, prediction accuracy also drops rapidly with RNA size (Mathews, Moss et al. 2010), further limiting its use. To find putative structures, or even evidence for the existence of stable structures, within the large datasets generated through modern tools other approaches are necessary.

One algorithm to address this is ScanFold (Andrews, Roche et al. 2018). ScanFold builds off of the Zuker algorithm, as implemented in the RNAfold program (Lorenz, Bernhart et al. 2011), however instead of trying to fold the entire sequence at once ScanFold takes a sliding window approach to find all local MFE structures across the long sequence. This is further enhanced by the calculation of a thermodynamic z-score that measures unusual sequence-ordered thermodynamic stability of the RNA fragment. For each window/fragment the nucleotides are shuffled and refolded multiple times to determine the mean MFE of the scrambled sequence set. A z-score is then determined by comparing the native/ordered MFE vs the average of the randomized sequences, normalizing by the standard deviation of MFE values. This thermodynamic z-score metric was to be an informative measure of how structured a given RNA sequence is by accounting for the importance of evolved ordering of the sequence on stability (Clote, FerrÉ et al. 2005). A lower z-score implies structural stability beyond what would be expected from nucleotide content: i.e., evolution may be acting to order this sequence for a structure/function relationship. This propensity of the z-score has been used to develop non-coding RNA prediction algorithms (Washietl, Hofacker et al. 2005) to deduce functional RNA structures across an array of targets (Moss, Priore et al. 2011, Moss and Steitz 2013).

Beyond just providing window level models of MFE structures and z-scores, ScanFold uses the MFE models and z-scores to develop z-score weighted consensus structures across the scanned sequence. Essentially providing unique models comprised of recurring base pairs favored in low z-score windows (i.e., the most ordered and stable base pairs). To rank to what degree an individual base pair shows a propensity to be found in unusually sequence-ordered stable structures, each base pair (i, j) is assigned a coverage-normalized z-score by adding the z-scores of the windows it occurs within and normalizing by the number of windows containing the base i. As this method can apply just as well to unpaired bases as to base pairs, unpaired bases are represented as a base pairing to itself. Using this scheme, ScanFold not only weights all predicted pairs, but allows for the reconstruction and filtering of structures comprised of unique low-z-score base pairs (Andrews, Roche et al. 2018). Approximate cutoffs were established with “low z-score” (high confidence) base pairs having average z-scores ≤ −2; “moderately low” (medium confidence) base pairs having average z-scores between −2 and −1; and low confidence negative z-score base pairs having average z-scores below 0 but above −1. This approach allowed for the identification of highly structured and functionally interesting targets across a variety of target transcripts and species: e.g., humans (Rouse, O’Leary et al. 2022, Rouse, Tompkins et al. 2024), viruses (O’Leary, Van Tompkins et al. 2022, Peterson, O’Leary et al. 2022, Dabiri, Rouse et al. 2025) and plants (Eisenhauer, Dapour et al. 2025).

The first major revision of ScanFold was ScanFold2, which greatly increased the speed of the scanning step by replacing explicit shuffling and refolding with a deep learning model trained to predict the mean and standard deviation of the curve created by shuffling a given sequence. By contrast, the “Fold” step has been relatively untouched, making it the logical focus for the next step of development. Here we introduce ScanFold3, enhancing the Fold step through algorithmic and language optimizations, achieving better scaling and an increase in speed from 38 to 129-fold over 500 b to 128 kb sequences respectively.

## Materials & Methods

An overview of the ScanFold3 algorithm can be found in Figure 1. As in previous versions of ScanFold, the input sequence is split into overlapping windows which are folded via Zuker’s Algorithm as implemented in RNAfold (Lyngsø, Zuker et al. 1999, Gruber, Lorenz et al. 2008). To account for the influence of sequence content on free energy, a thermodynamic z-score (Clote, FerrÉ et al. 2005) is calculated in each window and used as the primary metric for predicting thermodynamic stability. In the first version of ScanFold this was calculated by explicitly shuffling and refolding the input sequence multiple times, to create a normal curve of free energies; the z-score for the input sequence was calculated by subtracting the free energy of said sequence from the mean of this curve and dividing the result by the standard deviation of the set of energies (Andrews, Roche et al. 2018) Equation 1. In ScanFold2 and ScanFold3, a machine learning model which has been trained on data generated through this method is used instead to predict the thermodynamic z-score (Andrews, Rouse et al. 2022). These stages are referred to as the *Scan* step of ScanFold (ScanFold-Scan).

**Figure 1.**
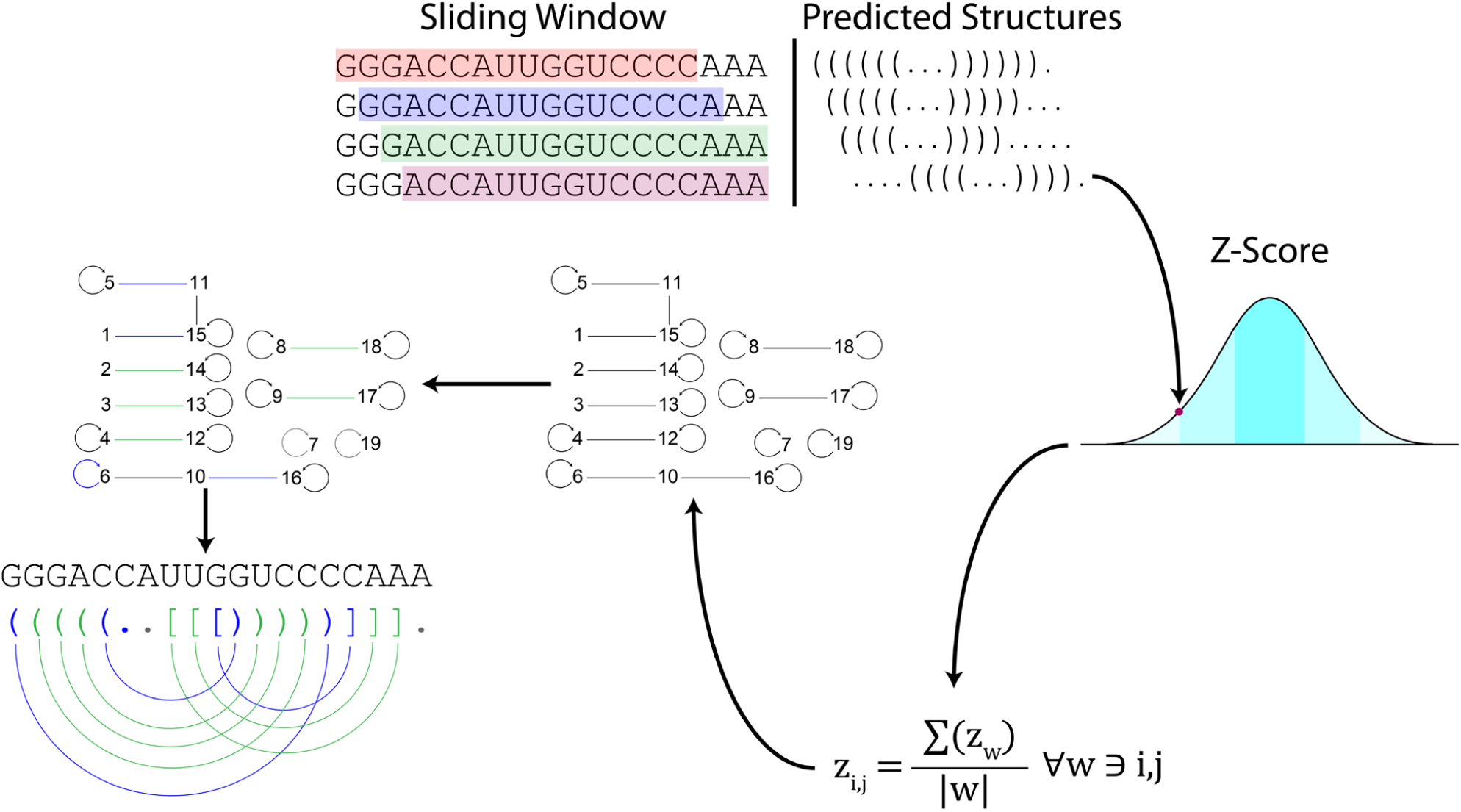
Overview of ScanFold3 algorithm. The input sequence is first split into overlapping windows, each of which is folded using Zuker’s Algorithm. Each base pair is assigned a normalized z-score derived from that of the windows it is found within (Equations 1, 2 & 3). Pairs are chosen by a greedy algorithm which keeps the most stable base pair in the event of conflicting base pairs to construct a final structure containing only non-competing base pairs.

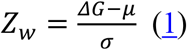

Following the *Scan* step is the *Fold* step (ScanFold-Fold), where z-score weighted structural data are collated across windows and used to construct a final consensus structure. Each base pair is assigned a normalized z-score, calculated by dividing the sum of the pair’s z-scores by its window coverage. This coverage represents the smallest of three values: the window length, the distance from the first nucleotide to the sequence start, or the distance from the second nucleotide to the sequence end. This is shown in Equation 2, where |w| is window size, s is sequence length, i is the first occurring nucleotide, and j is the second occurring nucleotide. Equation 3 shows the calculation for normalized z-score for a given base pair: the thermodynamic z-score of every window containing pair i, j are summed and divided by normalized window coverage. During this process, unpaired bases are represented as pairing to themselves, allowing the propensity of a base being unpaired across windows to be directly accounted for in the final model.

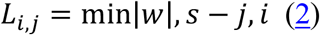

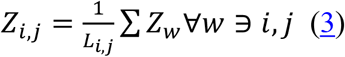

ScanFold3 builds off of previous versions of ScanFold by changing how a structure is built from these data. Previous versions of ScanFold mimicked the Scan step by iterating over each base and choosing the best partner for it to pair with after a short search for competing pairings for that target within 2 window lengths. ScanFold3 changes this by modeling this as a minimum perfect weighted matching problem on a graph where bases are modeled as nodes and pairs as edges. The minimum perfect weighted matching problem is the problem of finding the set of non-adjacent edges for a graph such that each node is connected to an edge with the lowest net weight. Unpaired bases are modeled as a self-edge; in order to use algorithms that do not allow self-edges, self-edges are removed, the graph is copied without self-edges, and these edges are represented as edges between analogous nodes on each graph.

Three algorithms for solving this problem were tested: Edmonds’ Blossom Algorithm (Edmonds 1965) and the Successive Shortest Path (SSP) Algorithm (Iri 1960), and a Greedy Approximation Algorithm (Figure 2). Edmonds’ Blossom was implemented using the Maximum Weighted Matching library distributed through Boost C++ Libraries (Ji and Rantwijk 2018), while the SSP Algorithm was coded in-house. While it does not necessarily find an optimal solution if one exists, the Greedy Approximation Algorithm using Introsort (MUSSER 1997) was considered as an alternative to finding the optimal solution due to better scaling and because it has been shown that greedy algorithms perform closer to optimal the sparser a graph is (Aronson, Frieze et al. 1998), and it was observed when testing the other algorithms that ScanFold graphs tend to be sparse (that is, the number of edges is not significantly higher than the number of nodes). It is worth noting that this algorithm does not explicitly represent base pairs as a graph, reducing the amount of overhead necessary.

**Figure 2.**
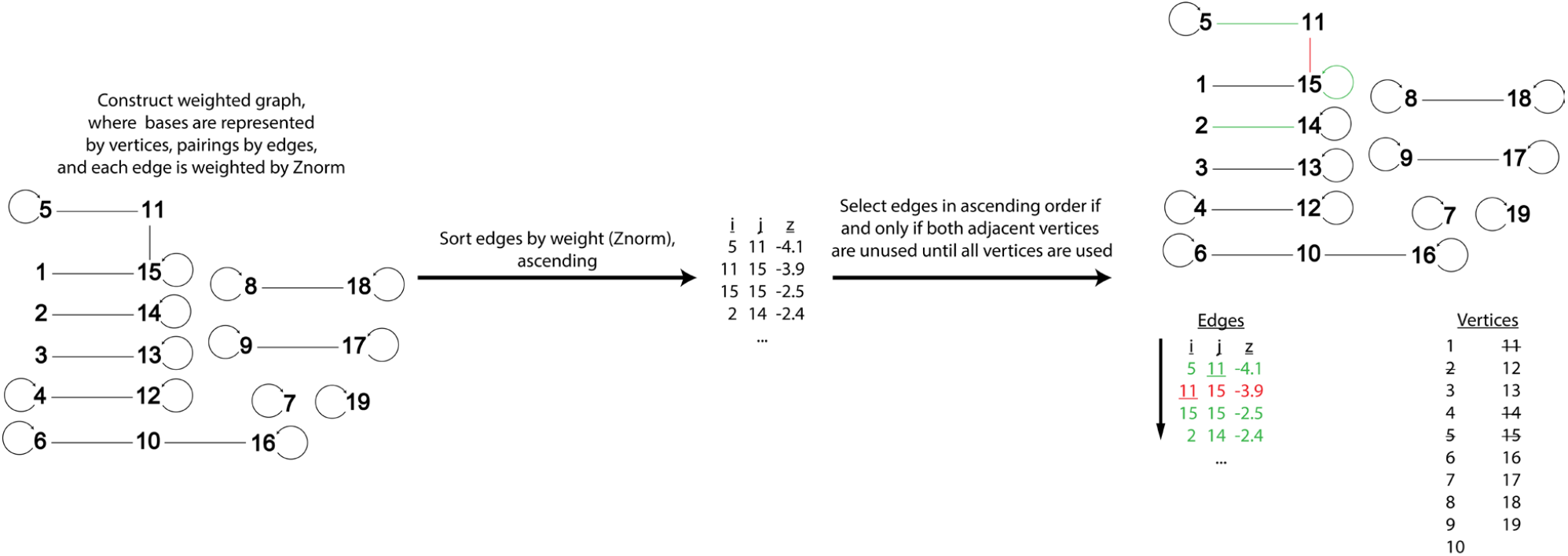
ScanFold3 Fold algorithm. Potential pairs are sorted from lowest (most stable) to highest (least stable) z-score, with unpaired nucleotides represented as self-pairing. Edges which do not compete with previously chosen edges are chosen for the final structure until no pairs remain or all bases have been paired.

In addition, ScanFold3 rewrites time-consuming portions of the code which had previously been written in Python in C++, which is on the order of ten times faster to run than Python on bioinformatics tasks (Fourment and Gillings 2008).

Up-to-date code for ScanFold3 alongside instructions for usage can be downloaded from GitHub at: https://github.com/moss-lab/ScanFold3. ScanFold3 code with script and validation data to reproduce results presented here can be found on Zenodo: https://doi.org/10.5281/zenodo.21384308 and https://doi.org/10.5281/zenodo.21223234, respectively.

To evaluate whether ScanFold3 preserved the ability to identify biologically functional RNA structures within long transcript contexts, we repeated the blind snoRNA (small nucleolar RNA) recovery analysis previously applied in the RNAStructuromeDB study (Dapour, et al. 2026). Annotated snoRNA coordinates were intersected with their corresponding host pre-mRNA transcript isoforms to define snoRNA-containing regions. The full host transcript sequences were then scanned with ScanFold3 without providing snoRNA annotations as input. After ScanFold3 motif extraction, predicted structural motifs were compared back to the known snoRNA coordinates in transcript-relative space. A snoRNA was considered recovered when a predicted motif overlapped the annotated snoRNA region by more than 20 nucleotides. For each snoRNA–isoform overlap, the most negative overlapping motif z-score was retained, and results were summarized across isoforms by calculating the mean z-score, best z-score, and number of recovered isoform overlaps per snoRNA.

## Results

Previous ScanFold-Scan results from the RNAStructuromeDB (Andrews, Baber et al. 2017) for Epstein-Barr Virus 1 were retrieved and used to benchmark speed. To evaluate how algorithm performance scales with dataset size, we constructed a benchmark test set using progressive truncations of the sequence. Specifically, we extracted structural windows covering the targets, sampling between lengths of 500 and 1,000, every 200 bases between 4,000 and 8,000, and at doubling distance from 8,000 up to 128,000 bases. Only the step of ScanFold involving the folding algorithm itself was tested; formatting and writing output and reading in ScanFold-Scan data were excluded. To determine if the SSP and Blossom algorithms were sufficiently scalable, tests were performed on the speed dataset comparing these to ScanFold2 and ScanFold3 with the greedy approximation algorithm (Fig. S1). The SSP algorithm matched the speed of ScanFold2 but ran out of memory at only 4 kb, tested with up to 1 terabyte of RAM. The Blossom algorithm reached 32 kb before running out of memory but was far slower than ScanFold2. Due to these results, it was determined that neither algorithm was appropriate for ScanFold3.

ScanFold3 when run with the greedy approximation algorithm showed a significant improvement to speed and scaling in comparison to ScanFold2 (Figure 3). A 38-fold speedup was observed at 0.5 kb, scaling up to a 129-fold speedup at 128 kb. When run on human chromosome 21, ScanFold3 was capable of producing results in 18 days, demonstrating its capacity to perform chromosome scale analysis on the order of weeks (Fig. S3). Speed benchmarks were performed on a compute node with 4x Intel Xeon Gold 6152 CPUs and 3TB of DDR4 memory.

**Figure 3.**
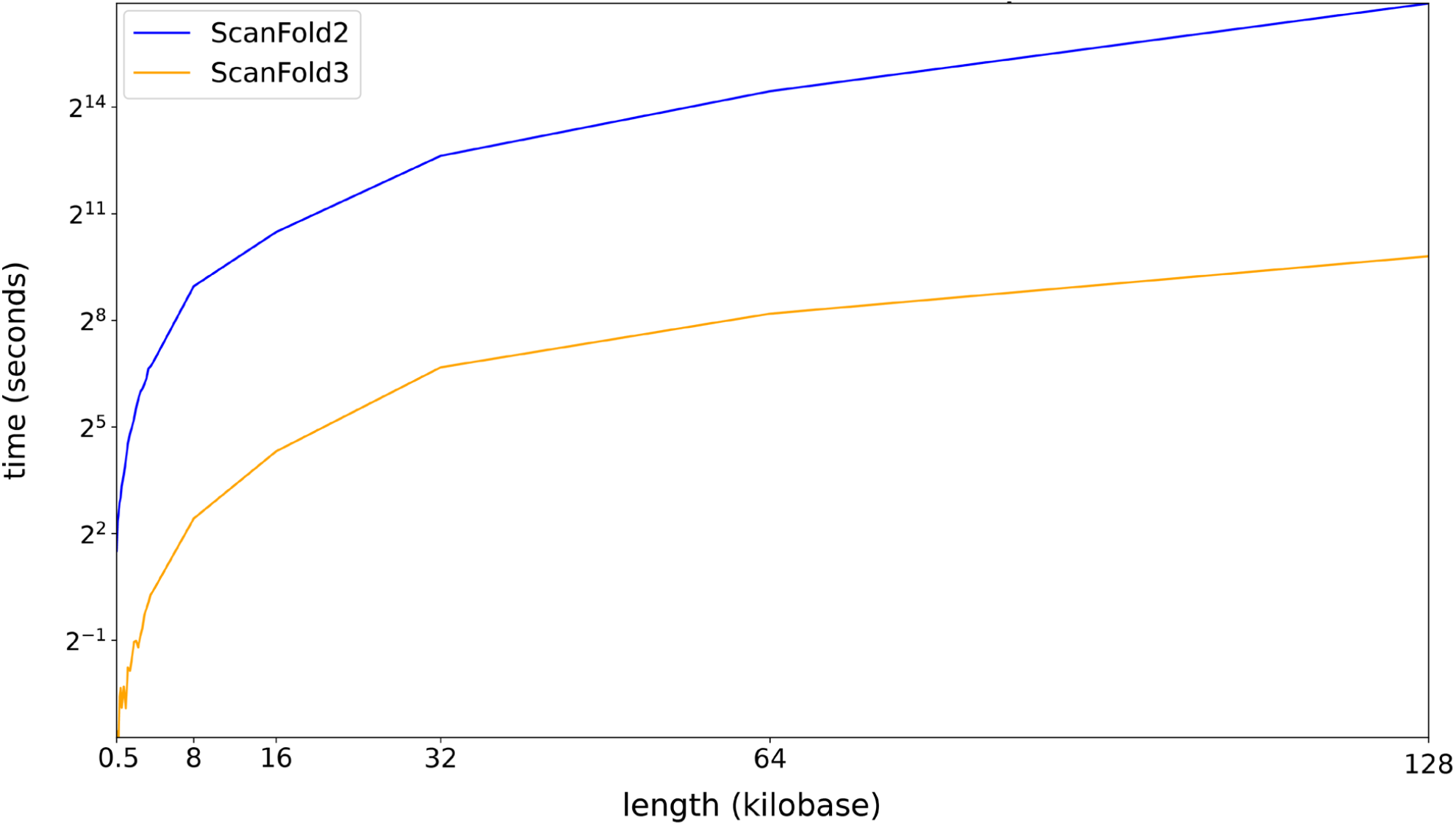
Comparison of speed between ScanFold2 (blue) and ScanFold3 (orange). Testing was performed on partitions of ScanFold-Scan data for Epstein-Barr Virus 1, taken from the RNAStructuromeDB. The difference in speed ranged from 38 times faster at 0.5 kb to 139 times faster at 128 kb. Datapoints were collected for every 100 bases between lengths of 500 and 1,000, every 200 bases between 4,000 and 8,000, and at exponential intervals scaled by powers of 2 from 8,192 up to approximately 128,000 bases.

To determine if ScanFold3 improved on the accuracy of ScanFold2, the bpRNA-1m dataset was retrieved on March 4, 2025 (Danaee, Rouches et al. 2018). 2,000 RNAs with lengths of at least 200 from this dataset were randomly selected. Of these, 1,403 were from the Comparative RNA Web (CRW) site (Cannone, Subramanian et al. 2002), 452 were from the RNA Families Database (Ontiveros-Palacios, Cooke et al. 2024), 78 were from the tmRDB (Zwieb, Gorodkin et al. 2003), 34 were from the Ribonuclease P Database (Brown 1998), 31 were from the Signal Recognition Particle Database (Larsen, Samuelsson et al. 1998), and two were from the Protein Data Bank (Berman, Henrick et al. 2007). These sequences had a mean length of 1031.7 nucleotides with a standard deviation of 594 nucleotides and a GC content of 53.6% with a standard deviation of 6.6% (File S2).

ScanFold3 was compared to ScanFold2 through several metrics: the positive predictive value (PPV; also known as precision), which is calculated from the ratio of true positives to true positives and false positives; the recall/sensitivity, which is the ratio of true positives to all positives; and the F1 score, which is calculated from the harmonic mean of PPV and recall. The complete output data generated from running ScanFold2 and ScanFold3 on these evaluation sequences is provided in File S1. This was performed both on the base pairs produced directly by ScanFold2 and ScanFold3, as well as the results after using these pairs as constraints for refolding the entire sequence with RNAfold (Gruber, Lorenz et al. 2008). Because the latter represents the recommended workflow for ScanFold and the method by which individual motifs are generated, it provides the most meaningful benchmark for assessing performance improvements.

The results are shown in Figure 4. When evaluated as an isolated predictor, ScanFold3 typically achieves higher PPV scores (Figure 4A) and lower recall scores (Figure 4B) compared to ScanFold2 across varying z-score filter thresholds. This high-precision, lower-sensitivity tradeoff is highly desirable for ScanFold, as it is meant to be used as a first pass to identify regions of interest for additional analyses: for example, to use with other folding algorithms like RNAfold (this functionality is included within ScanFold through the *--global_refold* flag). False negatives can be detected later in the pipeline, whereas false positives can contaminate later predictions if used as constraints. To determine whether this tradeoff improves predictions in this wider context, the structures derived from refolding using RNAfold guided by ScanFold constraints were analyzed through the same method. It was observed that at a given z-score filter there was no appreciable difference between ScanFold2 and ScanFold3 (Fig. S2).

**Figure 4.**
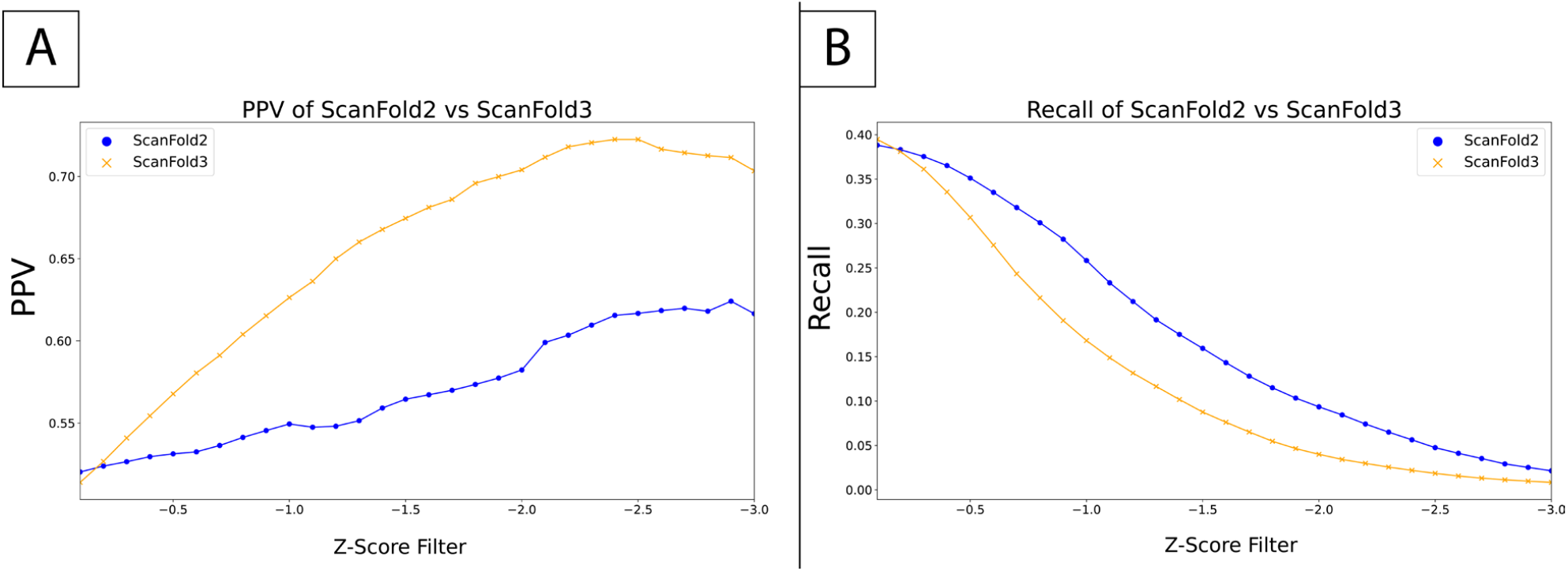
Comparison of predicted structures between ScanFold2 and ScanFold3 with a greedy approximation algorithm at a window size of 120. Benchmarking performed using structured RNAs from bpRNA-1m. A. Positive predictive value (PPV) of ScanFold2 (blue) and ScanFold3 (orange) structures when filtering out all pairs above a given z-score cutoff. B. Recall (sensitivity) of ScanFold2 (blue) and ScanFold3 (orange) when filtering out all pairs above a given z-score cutoff.

Previous work with ScanFold (Andrews, Roche et al. 2018) showed that a window size of 120 nt produced the highly reliable results. However, as the z-score filter level showed a change in performance between versions of ScanFold, it was possible that the same would be true of window size. To test this, a range of window sizes were tested at a filter level of −2, as shown in Figure 5. The F1 score alone suggests that performance improves as window size increases, however past a window size of 80 the instances of false positives steadily increases. Furthermore, lower window sizes are desirable as higher window sizes hamper performance in the Scan stage. 120 remains as an ideal compromise to keep the number of false positives down despite sacrificing performance in other metrics, particularly because extracted structures are refolded with RNAFold under default conditions. Notably, the average length structured RNAs archived within the RNACentral database falls below this cutoff. While long-range RNA base pairing interactions do play important roles in biology, many ncRNA families such as snRNA, Y RNA, and snoRNA fall within this range (Boivin, Faucher-Giguère et al. 2019). Notably, larger RNAs, while containing longer-range base pairs, are highly branched (Borodavka, Singaram et al. 2016), allowing ScanFold to detect the local structures that predominate the overall folding.

**Figure 5.**
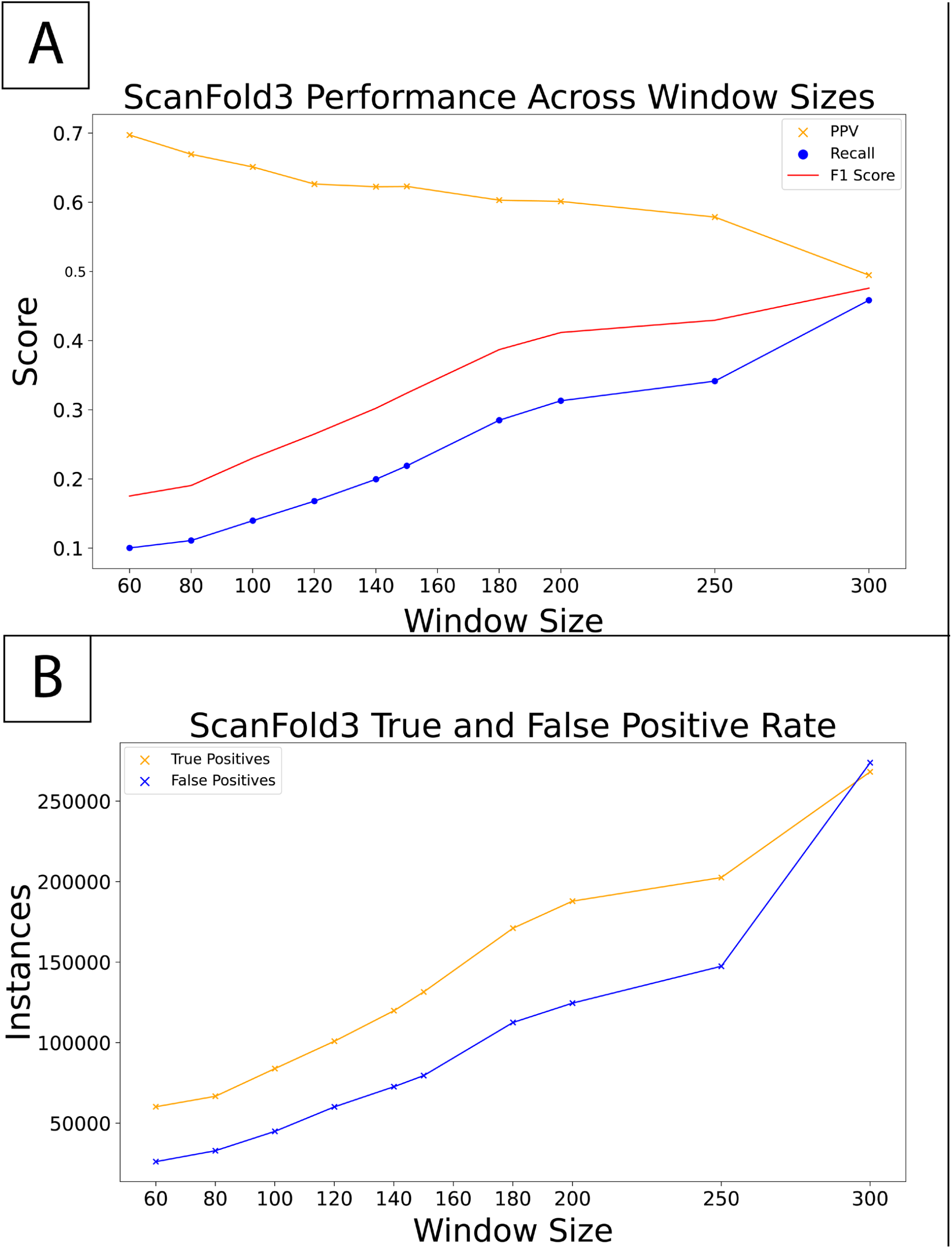
Performance of ScanFold3 across window sizes, on a set of reference sequences from bpRNA-1m. A. F1 Score (red), positive predictive value (orange), and recall (blue) of ScanFold3 across window sizes ranging from 60 to 300. B. Comparison of true positives (orange) and false positives (blue) across window sizes from 60 to 300. ScanFold3 uses a window size of 120 to maintain the speed of the Scan stage while minimizing the number of false positives which are sent forwards to be refolded.

To further evaluate whether ScanFold3 preserves biologically meaningful structural discovery in long transcript contexts, we repeated the blind snoRNA recovery analysis previously applied to the RNAStructuromeDB pre-mRNA dataset (Dapour, et al. 2026). In this analysis, full human transcript isoforms were scanned without providing snoRNA annotations to ScanFold3; annotated snoRNA coordinates were intersected with predicted structural motifs only after folding. Thus, snoRNAs served as embedded functional structural landmarks within the surrounding intronic background. ScanFold3 recovered 396 unique snoRNAs, compared with 335 recovered in the previous ScanFold2-based analysis (Dapour, et al. 2026), indicating improved recovery of annotated intronic structural elements. The recovered snoRNAs retained strong thermodynamic stability, with a median mean z-score of −3.13 (Figure 6). Furthermore, 82.1% of recovered snoRNAs (325/396) had mean z-scores ≤ −2.0, and 96.7% (383/396) had mean z-scores ≤ −1.0. These results demonstrate that ScanFold3 maintains the ability to distinguish evolved functional RNA structures from intronic background while increasing recovery of known snoRNA structural elements in full transcript context.

**Figure 6.**
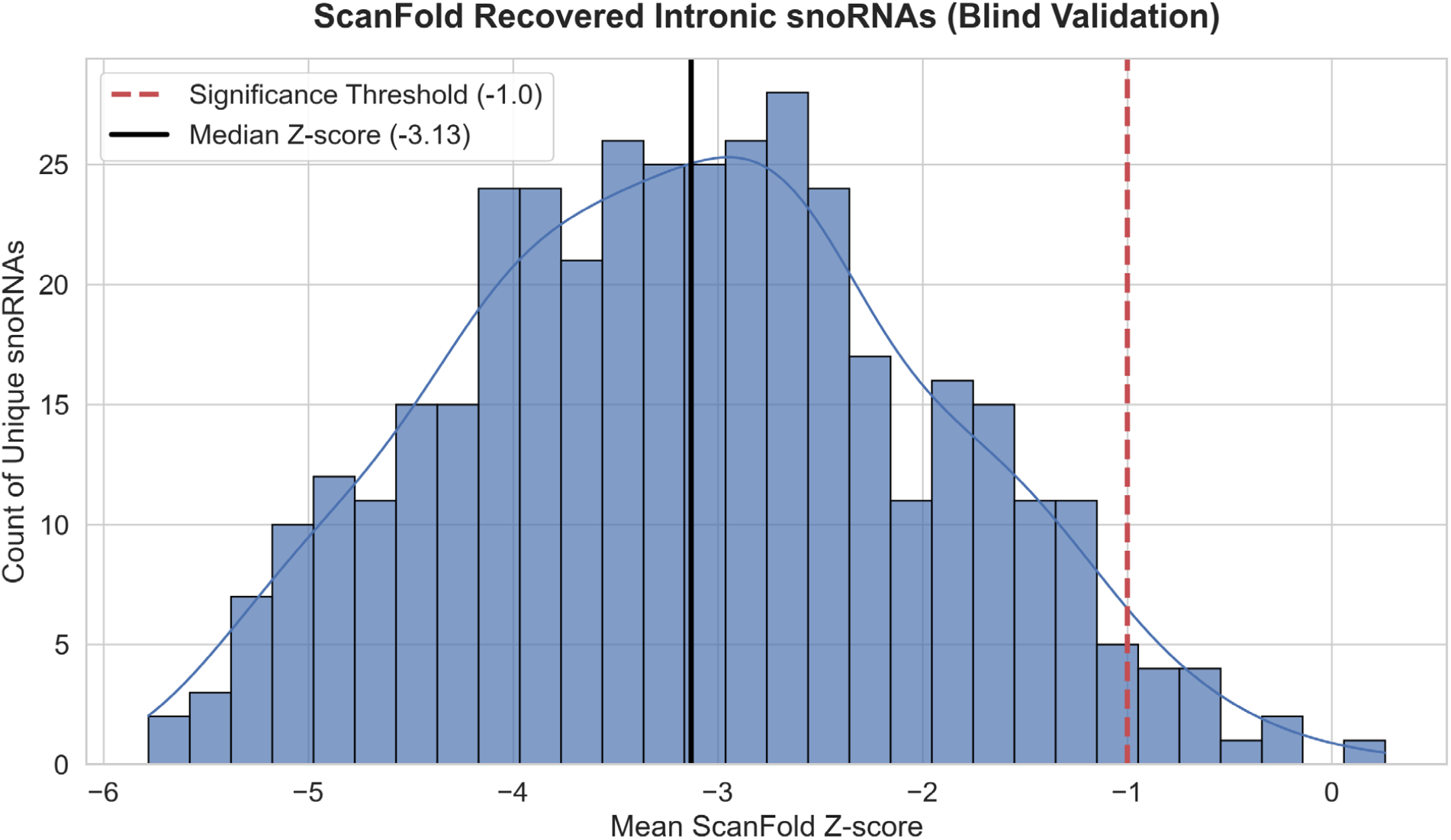
Blind recovery of intronic snoRNAs using ScanFold3. Histogram showing the distribution of mean ScanFold3 z-scores for 396 unique snoRNAs recovered from full host pre-mRNA transcript isoforms. Annotated snoRNAs were not provided to ScanFold3 during prediction; instead, predicted structural motifs were intersected with snoRNA coordinates after folding. The red dashed line indicates the z-score threshold of −1.0, and the black solid line indicates the median mean z-score of −3.13. Most recovered snoRNAs showed strong ordered thermodynamic stability, with 96.7% scoring ≤ −1.0 and 82.1% scoring ≤ −2.0, supporting the ability of ScanFold3 to recover functional structural elements from intronic background sequence.

## Discussion

ScanFold3 represents a significant enhancement in the characterization of RNA structure and thermodynamics: one that prioritizes the scalability required for modern genomic and transcriptomic analyses. By achieving up to a 139-fold increase in speed during the Fold stage for large sequences, ScanFold3 effectively removes the computational bottleneck that previously limited “omics” analyses. While ScanFold2 remains a robust tool, the transition to a C++ implemented greedy approximation algorithm in ScanFold3 allows researchers to quickly process datasets up to chromosome scale. Notably, when ScanFold3 is used in its primary role—generating structural constraints for global refolding—it shows no loss in performance for this increase in speed across all metrics, including F1 score and PPV. This confirms that the algorithmic optimizations do not sacrifice likely biological importance; rather, they optimize the identification of the most ordered/stable (likely evolutionarily conserved) structural motifs.

ScanFold has been designed for ease of installation and use, producing data in a wide variety of formats used in bioinformatics (Figure 7). This facilitates its incorporation into pre-existing bioinformatics pipelines, e.g., cm-builder, RNA Framework, R-Scape, etc. (Rivas, Incarnato, Morandi et al. 2018, Rivas, Clements et al. 2020, Peterson, O’Leary et al. 2023). Notably, ScanFold (through its RNAfold folding module) is able to directly incorporate chemical mapping constraints, e.g., from SHAPE or DMS probing; thus, it may also be used with (and enhance) various experimental analysis pipelines: such as SHAPE-MaP (Smola, Rice et al. 2015) and DMS-MaPseq (Zubradt, Gupta et al. 2017).

**Figure 7.**
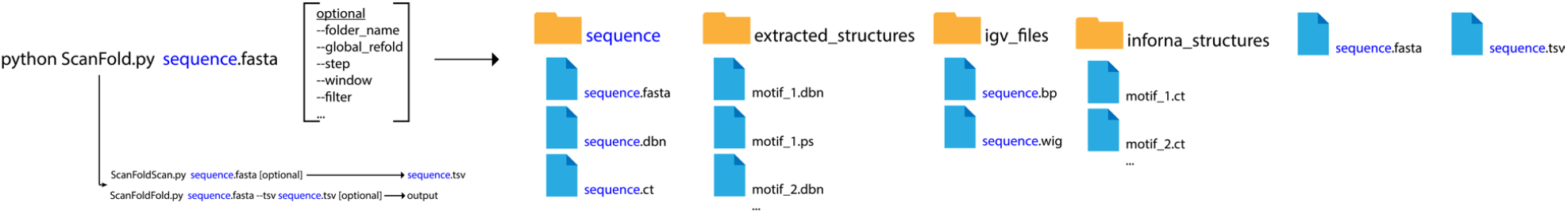
Graphical description of usage for ScanFold3. Detailed instructions can be found in the manual under the ScanFold3 github page. ScanFold is called through the ScanFold.py python script. A fasta file is necessary to run ScanFold; multiple sequences can be represented within the fasta file. Optionally, a specific output folder can be set, as can settings such as window size, whether to perform a global refold, and so on. ScanFold is split into two programs which automatically run back to back, ScanFoldScan and ScanFoldFold. ScanFoldScan produces a .tsv file containing all scanned windows, ScanFoldFold produces files containing sets of basepairs which can be used in other programs. Extracted_structures contains distinct motifs in the input sequence which have been extracted and refolded. Igv_files and informa_structures contain files meant to be used in the respective programs.

### Limitations

Despite these enhancements in function, some limitations remain with ScanFold3. Similar to previous ScanFold versions (and most folding algorithms) reliance on the Turner Rules for RNA folding prevents ScanFold3 from directly accounting for non-canonical base pairs and pairs involving naturally modified bases (e.g., inosine). Future work on ScanFold will explore methods for direct inclusion of non-canonical pairing information or nearest neighbor thermodynamics for modified bases (Hudson, Bloomingdale et al. 2013, Richardson and Znosko 2016, Wright, Force et al. 2018). Another limitation, common to all sliding window analyses, is the inability of ScanFold3 to directly consider longer-range base pairing (beyond the window sized used); it should be noted, however, that defining small, local structures, is extremely helpful in deducing the overall global architecture of a long RNA: e.g., by defining what nucleotides are *not* likely to form long-range pairs and which regions may be prone loop-loop interactions.

A final limitation of ScanFold3 to note is its inability to consistently predict pseudoknotted base pairs. Although the underlying folding algorithm used in ScanFold does not allow for pseudoknots, as scanning windows move through nucleotides, some are predicted to cross each other: thus, “knotted” base pairs and their thermodynamic parameters are stored within the log files of the program. Future work will explore the utility of these base pairs in pseudoknot prediction, as well as the inclusion of novel folding algorithms that can directly query pseudoknot folding: e.g., MXfold2 (Sato, Akiyama et al. 2021) or other AI/ML enhanced algorithms (Sacco, Bussi et al. 2025)

## Conclusions

ScanFold3 provides a fast and easy method for characterizing RNA thermodynamics, providing users with both regions within large sequences with a propensity for thermodynamic stability and a set of base pairs therein with a statistically significant propensity for stability. It is intended to be a first step in high-throughput bioinformatic pipelines (Peterson, O’Leary et al. 2023), capable of scaling to the ever increasing volume of sequenced transcriptomes and paring down these data to smaller structures amenable to other methods of analysis. Few other programs address the problem of quickly identifying promising regions in poorly or entirely un-annotated sequence data, thus ScanFold remains a valuable tool for best leveraging the most cutting-edge RNA folding techniques. As transcriptomic data continues to expand, ScanFold3 represents an important tool for narrowing the search for functional RNA structures across genomes and transcriptomes. This is particularly significant given the rising interest in RNA therapeutics (Tong, Childs-Disney et al. 2024) and, importantly, in identification of drug-targets within structured RNAs (Rouse, Andrews et al. 2022, Tong, Lee et al. 2023, Tong, Taghavi et al. 2025).

## Figures

**Supplemental Figure 1.**
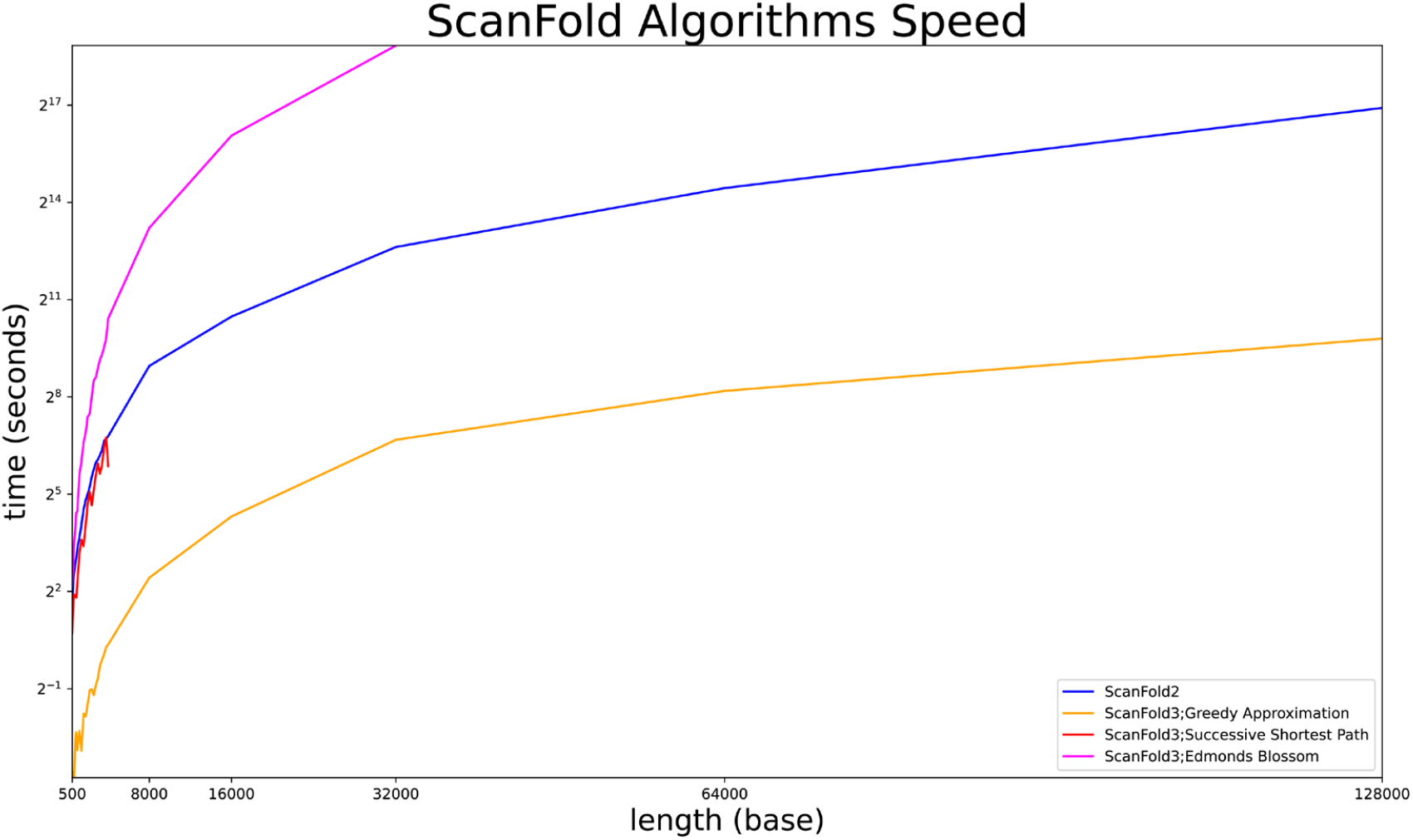
Comparison of speed of folding algorithms for ScanFold3, as well as ScanFold2. Blue: ScanFold2, run with default settings. Orange: ScanFold3 using a greedy approximation algorithm. Red: ScanFold3 using a Successive Shortest Path algorithm. Pink: ScanFold3 using Edmond’s Blossom algorithm. Early stopping indicates exceeding allocated memory. Testing was performed on partitions of ScanFold-Scan data for Epstein-Barr Virus 1, taken from the RNAStructuromeDB.

**Supplemental Figure 2.**
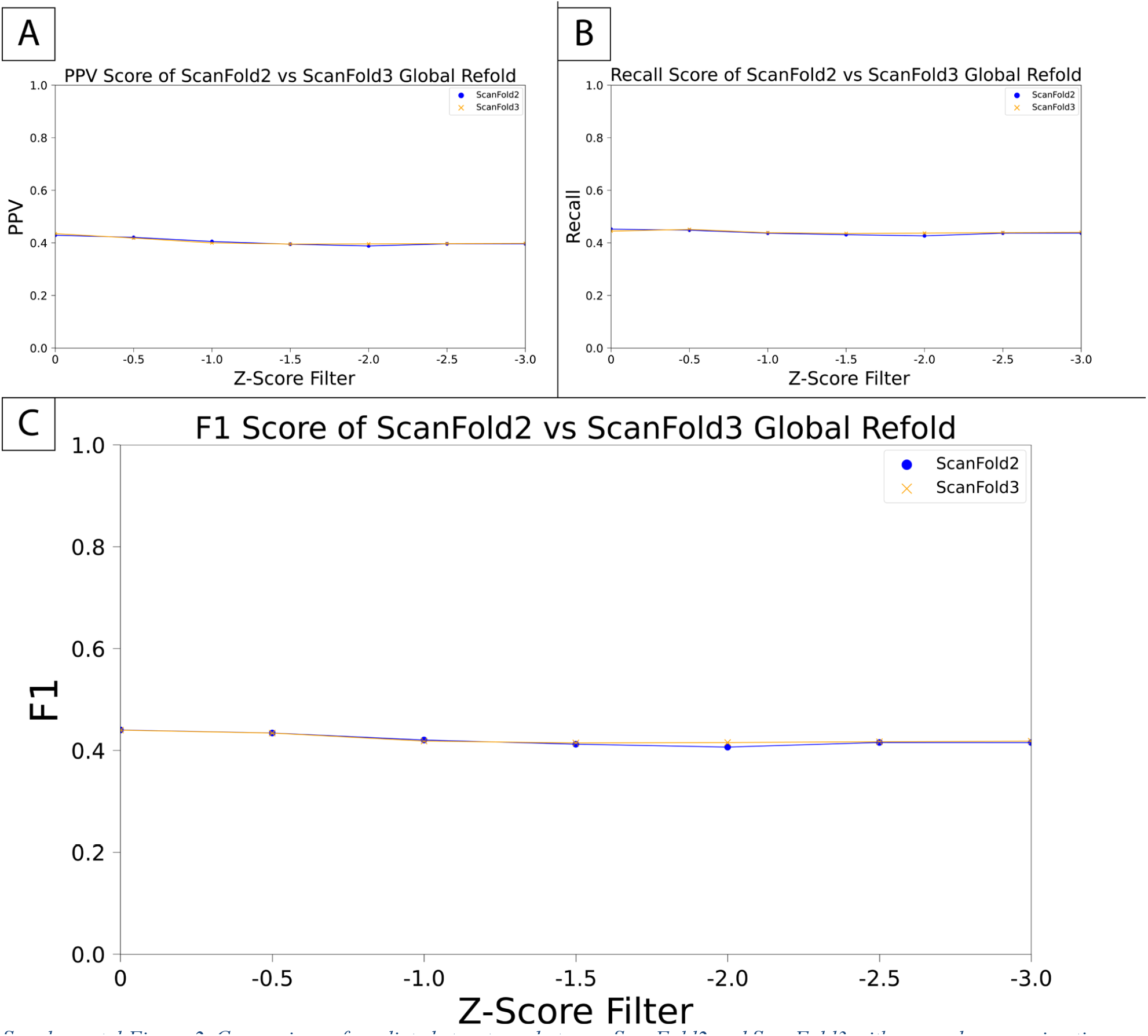
Comparison of predicted structures between ScanFold2 and ScanFold3 with a greedy approximation algorithm at a window size of 120 after performing a global refold with RNAFold using pairs with z-scores below filter levels (x-axis) as constraints. Benchmarking performed using structured RNAs from bpRNA-1m. A. PPV of ScanFold2 (blue) and ScanFold3 (orange) when pairs below a predicted cutoff were used as constraints for RNAFold. B. Recall of ScanFold2 (blue) and ScanFold3 (orange) when pairs below a predicted cutoff were used as constraints for RNAFold. C. F1 Score (harmonic mean of PPV and recall) of ScanFold2 (blue) and ScanFold3 (orange) when pairs below a predicted cutoff were used as constraints for RNAFold. ScanFold3 alone shows better PPV but lower recall than ScanFold2. When used as a constraint for RNAFold these differences vanish and neither performs better than the other.

**Supplemental Figure 3.**
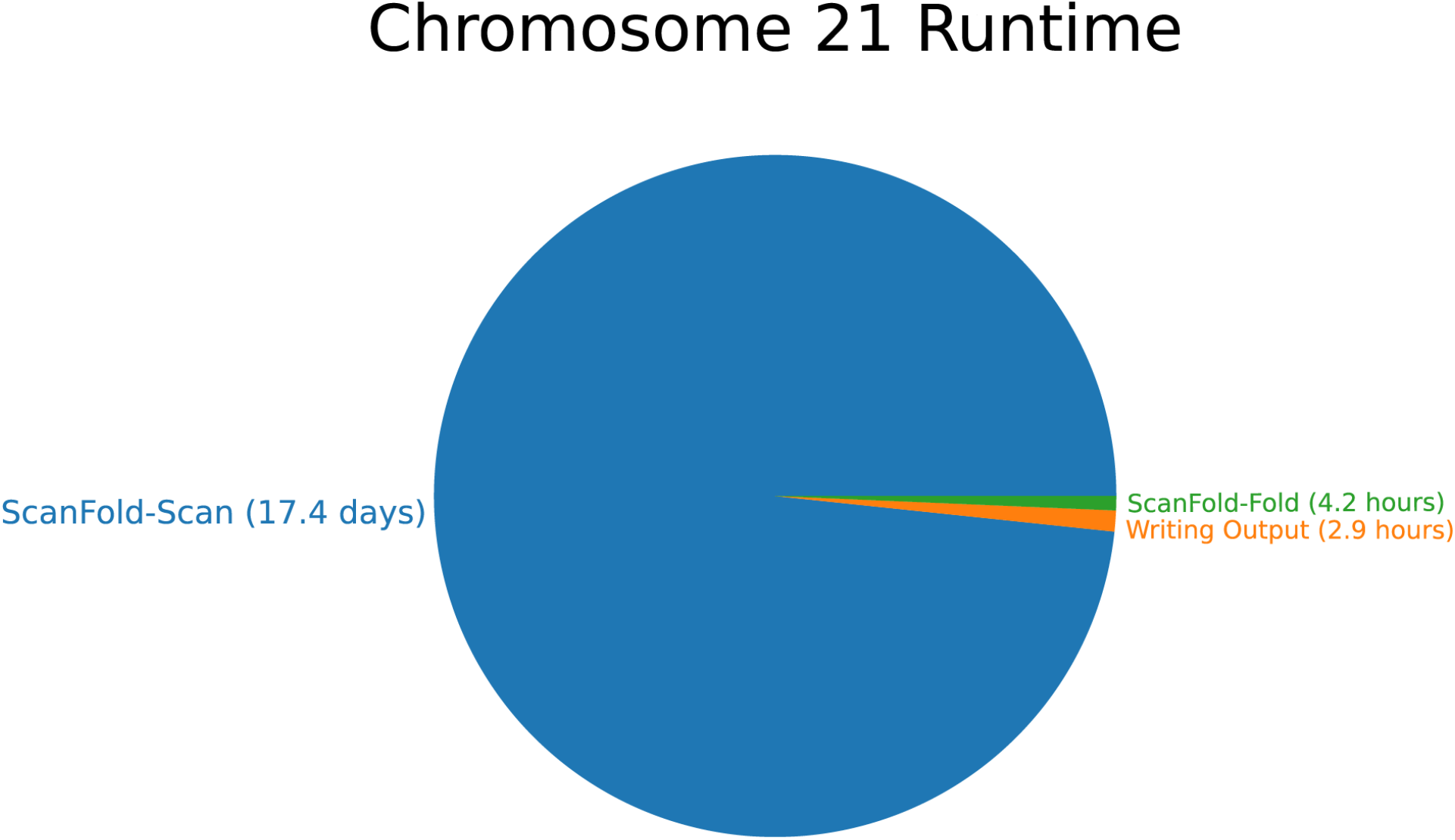
Speed benchmark on chromosome 21. Run on a node with 4x Intel Xeon Gold 6152 CPUs and 3TB of DDR4 memory. The Scan stage completed in 1,504,327 seconds (approximately 17.4 days). The Fold stage completed in 15,165 seconds (approximately 4.2 hours). Writing output from the Fold stage was completed in 10,299 seconds (approximately 2.9 hours)

## Acknowledgements

Thank you to the Iowa State University Research IT group for their support over the course of this project and members of the Moss Lab for their input.

